# Modification and validation of a reference real-time RT-PCR method for the detection of a new African horse sickness virus variant

**DOI:** 10.1101/2025.10.06.680186

**Authors:** Jorge Morales, María José Ruano, Cristina Tena-Tomás, Antoinette Van Schalkwyk, Eleni-Anna Loundras, Marta Valero-Lorenzo, Ana López-Herranz, Marco Romito, Carrie Batten, Rubén Villalba, Montserrat Agüero

## Abstract

African horse sickness (AHS) is a disease affecting equids caused by the AHS virus (AHSV). The World Organization for Animal Health (WOAH) includes AHS as a notifiable disease and, upon detection within the European Union, immediate control and eradication measures are mandated. Thus, validated diagnostic methods for rapid AHSV detection are essential. The Agüero 2008 and Guthrie 2013 rRT-PCR methods have been widely validated for detection of any AHSV strain and are included as reference rRT-PCRs in the WOAH manual. However, the WOAH reference laboratory for AHS in the Republic of South Africa (RSA), reported an AHSV variant undetected by the Agüero 2008 rRT-PCR. Therefore, a set of modified primers and probe, containing degenerate positions to avoid mismatches with the sequence of the new RSA strain, was developed. The modified-Agüero method was validated by the WOAH reference laboratories in Spain and the UK employing a broad collection of AHSV strains, clinical samples as well as a synthetic RNA mimicking the target sequence of the new RSA AHSV variant (AHSV-sRNA-RSA). Comparative assessment of the modified-Agüero versus the WOAH reference rRT-PCRs, showed that the modified method exhibited a good diagnostic performance and enabled the detection of the new RSA AHSV variant nucleic acid.

## Introduction

African horse sickness (AHS) is a viral disease that affects equids. It is caused by the African horse sickness virus (AHSV), an orbivirus from the *Sedoreoviridae* family, and transmission occurs through the bite of *Culicoides*, making environmental factors and midge population densities critical in the spread of the disease (Assefa et al., 2022; Mellor & Hamblin, 2004; Zientara et al., 2015). The fact that nine distinct serotypes of AHSV have been identified is significant for diagnostic and vaccination purposes (Coetzer & Tustin, 2004; European Union, 2016; King et al., 2020; Lu et al., 2020; Mellor & Hamblin, 2004; World Organisation for Animal Health, 2020, 2025b; Zientara et al., 2015). The variety of clinical presentations in AHS adds significant complexity to its management and control. The four forms—horse sickness fever, cardiac, mixed and pulmonary forms—differ in severity, with the pulmonary form being particularly devastating (Coetzer & Tustin, 2004; Mellor & Hamblin, 2004). The high mortality rate associated with the respiratory form, sometimes exceeding 95% in non-vaccinated animals, underscores the critical importance of vaccination in preventing outbreaks and reducing fatalities (Coetzer & Tustin, 2004; Mellor & Hamblin, 2004; Zientara et al., 2015). Besides the relevant impact of AHS in equid welfare, it is estimated that African horse sickness results in an annual economic loss of approximately $95 million USD in sub-Saharan Africa, where it is endemic (Redmond et al., 2022).

The geographical spread of AHS outside sub-Saharan Africa highlights the potential for the disease to affect equine populations globally. Historical outbreaks in the Middle East, North Africa, and later in Europe and Asia, show how quickly the virus can spread, especially when conditions are conducive for the vector to thrive (Mellor & Hamblin, 2004; Zientara et al., 2015). The outbreak caused by AHSV-4 in Spain, Portugal and Morocco in the late 1980s, and more recently in Southeast Asia by AHSV-1 (King et al., 2020; Lu et al., 2020; Mellor P.S., 1993; World Organisation for Animal Health, 2020), further exemplify the global risks posed by the disease to animal health and the equine industry worldwide. The rapid spread of AHS and its high mortality rate in unvaccinated horses emphasize the need for effective surveillance, vaccination programs, and vector control strategies, especially in non-endemic regions, to mitigate the impact of this serious disease (Castillo-Olivares, 2021; Dennis et al., 2019; Redmond et al., 2022),

The World Organisation for Animal Health (WOAH) includes AHS as a notifiable disease, reflecting the global importance of monitoring and controlling the disease (World Organisation for Animal Health, 2025b). Therefore, AHS must be reported by affected countries, enabling the WOAH to assist in coordination of global response efforts to prevent further spread of the disease (World Organisation for Animal Health, 2024). Within the European Union (EU), AHS is considered one of the animal diseases for which immediate control and eradication measures are mandated upon AHSV detection. This is included in the EU’s comprehensive “Animal Health Law,” which sets protocols for disease management, surveillance, and eradication, helping to prevent outbreaks that could impact both equine health and the agricultural economy in the EU (European Union, 2016). The international legislative frameworks emphasize the importance of vigilance, rapid response, and collaboration to minimize the impact of AHS and protect equine populations worldwide. To meet these goals, the availability of validated, effective and harmonised diagnostic methods is essential. In this regard, there are several real-time reverse-transcription polymerase chain reaction (rRT-PCR) methods available for AHS serogroup detection (Agüero et al., 2008; Bachanek-Bankowska et al., 2014; Fernández-Pinero et al., 2009; Guthrie et al., 2013; Monaco et al., 2011; Rodriguez-Sanchez et al., 2008) and serotype identification (Bachanek-Bankowska et al., 2014; Koekemoer, 2008; Villalba et al., 2024; Weyer et al., 2015). Specifically, the rRT-PCR methods described by Agüero (Agüero et al., 2008) and Guthrie (Guthrie et al., 2013; Quan et al., 2010) are included as reference rRT-PCRs for AHSV detection in the WOAH manual (World Organisation for Animal Health, 2025a). Both methods target segment 7 of the viral genome, allowing the detection of any AHSV strain, regardless of the serotype. Currently, the Agüero 2008 rRT-PCR is widely used by EU national reference laboratories (EU-NRLs) for AHSV molecular diagnosis. In fact, 19 out of 27 (70.4%) EU-NRLs participating in the last proficiency test organized by the AHS EU Reference Laboratory (EURL), used the Agüero 2008 as rRT-PCR method for AHS serogroup detection (EURL, personal communication).

Recently, the Pretoria University (RSA) and the WOAH reference laboratory in the RSA at the Agricultural Research Council – Onderstepoort Veterinary Institute (ARC-OVI), reported to the WOAH and EURL for AHS, the Laboratorio Central de Veterinaria (LCV), based in Algete, Madrid (Spain), that the Agüero 2008 rRT-PCR was unable to detect a putatively novel AHSV strain from Western Cape. The sample submitted to the ARC-OVI was successfully amplified using the hemi-nested assay (Bremer et al., 1998) and typed as serotype 9 (van Schalkwyk et al 2019). Moreover, it was reported that this strain had been previously detected by the Guthrie 2013 rRT-PCR method.

To further underpin the lack of sensitivity of the Agüero 2008 rRT-PCR to detect the novel AHSV variant, the WOAH reference laboratory in RSA obtained the partial sequence of the segment 7 covering the region targeted by the Agüero 2008 and Guthrie 2013 rRT-PCR methods. Despite being a highly conserved region of the virus genome, some mismatches were identified between the primers and probes of both methods and the new strain target sequence. Consequently, modified primers and probe based on the Agüero 2008 method containing degenerate positions to avoid mismatches were designed, tested *in-silico* and validated. Given that the new AHSV strain detected in the RSA was not available at the WOAH Reference Laboratory in Spain, two synthetic RNAs containing the target sequence of the Agüero 2008 and Guthrie 2013 rRT-PCRs were designed: -one positive control mimicking the target sequence of the reference AHSV strains (AHSV-sRNA-C+), -and another with the segment-7 target sequence of the undetected RSA strain (AHSV-sRNA-RSA). These synthetic RNAs were used for the comparative assessment of the modified-Agüero rRT-PCR versus the WOAH manual reference methods, in addition to a wide variety of AHSV strains and clinical samples from the collections kept at the WOAH Reference Laboratories in the UK (The Pirbright Institute) and in Spain (EURL, LCV). In this study we present the optimization and validation of the modified-Agüero method, which allows the reliable detection of the new RSA AHSV strain, as well as the AHSV collection strains and clinical samples.

## Material and methods

### Nucleid Acid Extraction

200 μl of EDTA blood samples, tissue homogenates or viral suspensions were subjected to nucleic acid extraction using the BioSprint® 96 DNA Blood Kit (Qiagen, Hilden, Germany). Nucleic acids were eluted in a final volume of 50 μl of nuclease-free water and kept at −80ºC until used.

### Primers and probe design

The primer and probe sequences of the original Agüero method (Agüero et al., 2008) were modified to target the RSA strain sequence provided by the RSA WOAH reference lab. Subsequently, in the reverse primer, AHS-R-A2024 (5’- CTAATGAAAGCGG**Y**GACCGT-3’), a degenerate nucleotide was introduced in the 14^th^ position, whereas in the probe, AHS-R-A2024 (FAM-GCTAGC**R**GC**Y**TACCACTA-MGB), degenerate nucleotides were included in the 7^th^ and 10^th^ positions. Additionally, these oligonucleotides as well as the ones of the reference RT-PCR methods were aligned with sequences encoding the AHSV VP7 currently available in the GenBank database (n=227) using ClustalW 1.8 (Thompson et al., 1994) to detect mismatches that may compromise AHSV detection.

### Design of the synthetic AHSV RNAs

A synthetic RNA containing the sequence of the undetected South African strain targeted by the Agüero 2008 and Guthrie 2013 rRT-PCR methods plus 21 nucleotides on each end of the molecule (AHSV-sRNA-RSA) was designed and ordered. Likewise, a synthetic RNA equalling the target sequence shared by most AHSV reference strains (AHSV-sRNA-C+) was designed as a positive control. These nucleic acids, of 120 nulcelotides in length each, were synthesized by Metabion International AG (sup. Figure 1). Upon arrival, 1 nmol of each dry synthetic RNA pellet was used to prepare specific solutions containing 10^9^ molecules/μl of each synthetic RNA in water and kept at −80ºC until further use. Ten-fold serial dilutions of these synthetic RNAs solutions were used for the comparative assessment of the diagnostic performance of the modified-Agüero method versus the WOAH reference rRT-PCR methods.

**Figure 1.**
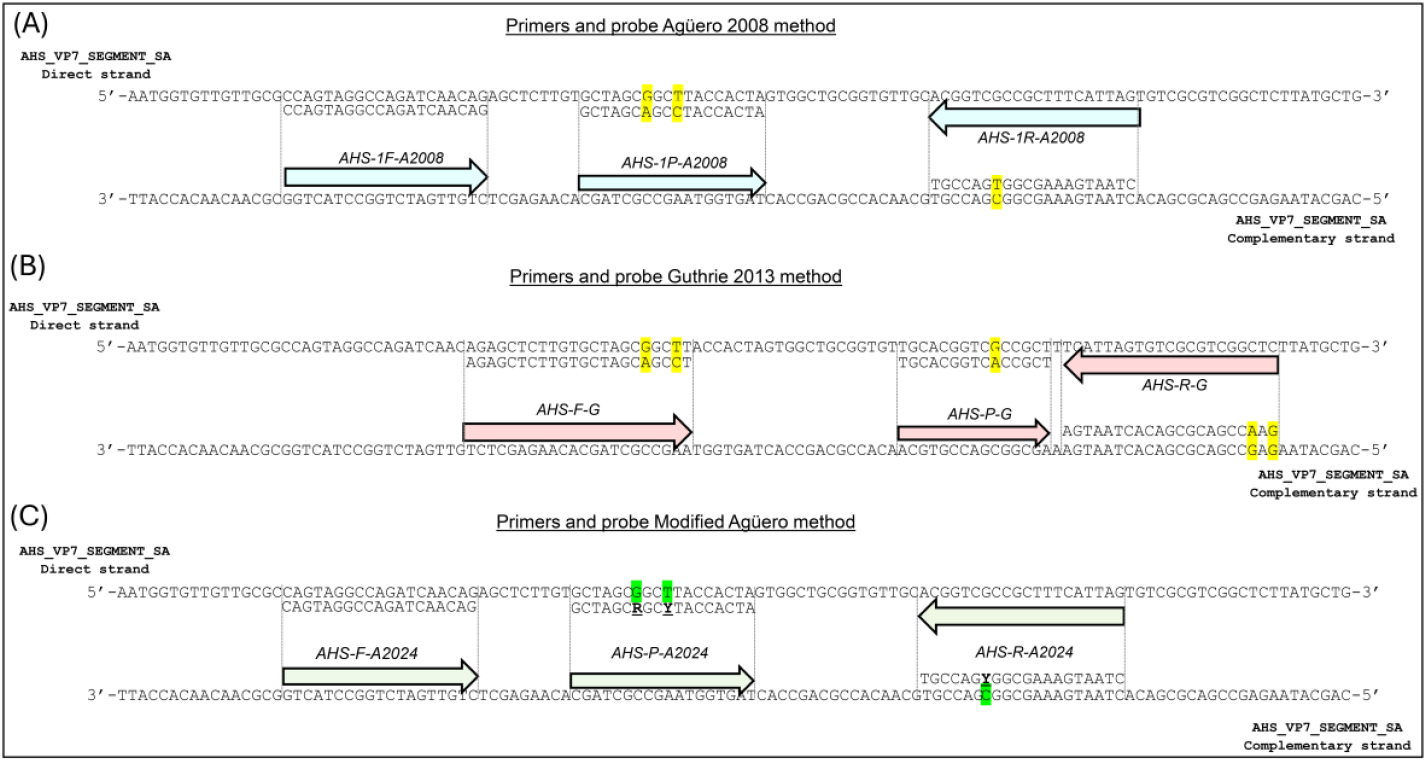
Alignment of the primers/probe described by Agüero (2008), Guthrie (2013) and the modified Agüero method with the segment 7 of the new AHSV strain from RSA. (A) Alignment of the forward (AHS-1F-A2008) and reverse (AHS-1R-A2008) primers and probe (AHS-1P-A2008) of the Agüero 2008 method with the target sequence of the AHSV strain from RSA. (B) Alignment of the forward (AHS-F-G) and reverse (AHS-R-G) primers and probe (AHS-P-G) of the Guthrie 2013 method with the target sequence of the AHSV strain from RSA. (C) Alignment of the forward (AHS-1F-A2024) and reverse (AHS-1R-A2024) primers and probe (AHS-P-A2024) of the modified-Agüero rRT-PCR method with the target sequence of the AHSV strain from RSA. The sequences are shown as cDNA, the primers and probes are depicted by coloured arrows and located at the annealing site of the cDNA target strand. The sequence of the primers and probes are shown aligned with the opposite strand to facilitate the visualisation of mismatches. Mismatches are highlighted in yellow and modifications of the primer/probe are shown in green.

### Serogroup-specific real-time RT-PCR

To test the performance of the modified primer and probe, rRT-PCR assays were carried out in the same conditions as the original method following the rRT-PCR protocol described in (Durán-Ferrer et al., 2022). Briefly, rRT-PCR was performed in a final volume of 20 μl using the commercial kit AgPath-ID™ One-Step RT-PCR Reagents (Applied BioSystems, Whaltman, MA, USA). Samples were classified as positive when a typical amplification curve was obtained and the Ct value was lower or equal to 35 within 40 PCR cycles (Ct ≤ 35), inconclusive when 35 < Ct < 40, and negative when a Ct > 40 or no Ct was obtained.

### Viral strains and clinical samples

The nine AHSV reference strains (serotypes 1 to 9) and thirty-four additional strains from the EURL’s AHSV reference collection were used for the validation of the modified-Agüero rRT-PCR. Further, eleven AHSV strains from The Pirbright Institute presenting mismatches with the Agüero 2008 and Guthrie 2013 primers and/or probe were included in this validation to evaluate the inclusivity of the modified method. To assess the specificity (exclusivity), representative strains of various orbiviruses were employed, including reference strains of bluetongue virus (BTV) corresponding to notifiable serotypes 1 to 24, epizootic haemorrhagic disease virus (EHDV) serotypes 1, 2, 4, 5, 6, 7, and 8, as well as equine encephalosis virus (EEV) serotype 3. Additionally, viruses known to induce disease in equids, such as West Nile virus lineage 1 and equine herpesvirus serotypes 1 and 4, were incorporated (sup. Table 1). All orbivirus strains used in this investigation from the EURL collection were propagated in BHK-21 clone 13 cells (American Type Culture Collection, ATCC-CCL-10TM), Vero cell monolayers (ATCC-CCL81), or KC cells derived from *Culicoides sonorensis* (Wechsler & Mcholland, 1988). Control rRT-PCR analyses included uninfected cell lines. Clinical samples of equines were used to assess the diagnostic parameters, which included: 30 EDTA blood and 25 tissue samples from the 2015-2017 outbreaks in Kenya, which is an AHS endemic country, 24 spleen samples from the AHS outbreak in Spain (1987–1990), and 28 EDTA blood samples corresponding to horses from Spain obtained between 2019– 2022 (AHS-free status country). Tissue samples were collected post-necropsy and preserved by storage at ultra-low temperatures (−80ºC). Viral suspensions were prepared through serial passages in cell cultures (KC, Vero, or BHK-21), followed by clarification via centrifugation at 805 x g for 10 minutes, and subsequently stored at −80ºC. EDTA blood samples were maintained at refrigerated conditions between 2 and 8ºC or stored at -80ºC until laboratory analysis.

**Table 1.** AHSV segment 7 sequences from GenBank (2009 – 2023) showing mismatches with primers and probes.

| Pattern | Number of sequences | GenBank accession number (AN) | Description of mismatches with the Agüero 2008 primers and probe |  |  |
| --- | --- | --- | --- | --- | --- |
|  |  |  | Position-nucleotide change from 5' |  |  |
|  |  |  | Forward | Probe | Reverse |
| A1 | 17 | KT715637.1 KT715607.1 KT030586.1<br>KT030616.1 KF860002.1 HM035402.1<br>HM035399.1 HM035396.1 KP939763.1<br>KY471520.1 KY471519.1 KP939655.1<br>KP939759.1 KP939758.1 KP939760.1<br>KP939761.1 KP939459.1 | 3-A>G |  |  |
| A2 | 1 | KT030396.1 | 11-A>G |  |  |
| A3 | 2 | HM035370.1 HM035365.1 | 12-G>A |  |  |
| A4 | 20 | KT030666.1 KF860042.1 KP009697.1<br>HM035403.1 HM035376.1 KP940201.1<br>HM035373.1 KP940190.1 KP940079.1<br>KP940078.1 KP940076.1 KP940075.1<br>KP940074.1 KP940073.1 KP940072.1<br>KP940071.1 KP939886.1 KP939879.1<br>KP939402.1 U90337.1 | 15-C>T |  |  |
| A5 | 20 | MT586218.1 KT030526.1 KT007174.1<br>KT187063.1 KT187033.1 KT030476.1<br>KT030466.1 KP009727.1 KP009687.1<br>KP009627.1 HM035392.1 HM035382.1<br>HM035378.1 HM035374.1 HM035364.1<br>HM035361.1 MT711964.1 KP939647.1<br>KP939537.1 KP939534.1 | 18-C>T |  |  |
| A6 | 1 | KP940080.1 | 15-C>T<br>6-A>T |  |  |
| A7 | 14 | KT030416.1 KT030556.1 KT030536.1<br>HM035389.1 HM035388.1 HM035383.1<br>HM035381.1 HM035377.1 HM035371.1<br>HM035363.1 KP940196.1 KP939976.1<br>KP939971.1 HM035393.1 | 18-C>T<br>3-A>G |  |  |
| A8 | 18 | HM035397.1 HM035390.1 HM035386.1<br>HM035385.1 HM035384.1 HM035380.1<br>HM035379.1 HM035368.1 HM035367.1<br>KP939975.1 KP939970.1 KP939762.1<br>KP939878.1 KP939882.1 KP939973.1<br>KP939888.1 KP939887.1 KP939884.1 | 9-C>T |  |  |
| A9 | 1 | KP939454.1 | 9-C>T |  | 8-A>G |
| A10 | 1 | KP939655.1 | 3-A>G | 10-C>T* |  |
| A11 | 9 | KP940199.1 KP940193.1 KP939657.1<br>KP939652.1 KP939650.1 KP939648.1<br>KP939536.1 KP939535.1 KP939456.1 |  | 10-C>T* |  |
| A12 | 2 | KP939767.1 KP939454.1 |  |  | 8-A>G |
| A13 | 3 | KT715617.1 KT030636.1 KP939881.1 |  |  | 4-A>G |
| A14 | 1 | KT030546.1 |  |  | 15-G>A |
| Pattern | Number of sequences | Genbank accession number (AN) | Description of mismatches with the Guthrie method primers and probes |  |  |
|  |  |  | Position-nucleotide change from 5' |  |  |
|  |  |  | Forward | Probe | Compl. Reverse |
| G1 | 10 | KP939650.1 KP939648.1 KP939456.1<br>KP939535.1 KP939536.1 KP939652.1<br>KP939657.1 KP939655.1 KP940193.1<br>KP940199.1 | 21-C>T |  |  |
| G2 | 2 | KT715617.1 KP939881.1 |  |  | 18-A>G |
| G3 | 1 | KT939762.1 |  |  | 5-C>T |

\*This mismatch was corrected by modifying the probe in the modified-Aguero method

### rRT-PCR validation parameters evaluated

Validation parameters of the modified-Agüero method were mainly assessed by comparison to the reference rRT-PCR method Agüero 2008. Analytical sensitivity was determined using ten-fold serial dilutions of the nine AHSV reference strains (serotypes 1 to 9). Analytical specificity (exclusivity) of the modified rRT-PCR method was evaluated by analysing other orbiviruses, several viruses affecting horses, and the most common cell lines used for AHSV propagation. Analytical specificity (inclusivity) was assessed using the EURL’s AHSV strains collection. Additionally, to further appraise inclusivity, several AHSV strains from The Pirbright Institute were analysed by using the three methods (Agüero 2008, Guthrie 2013 and modified-Agüero). The diagnostic performance of the assay was evaluated using clinical samples (EDTA blood and tissue samples) from the AHS outbreaks in Spain 1987-1990, the 2015-2017 outbreaks in Kenya, and horse blood samples from free-of-AHS areas. Finally, intra-assay repeatability was checked performing the analytical sensitivity tests in duplicate and calculating the Ct difference between assays.

### Statistical analysis

To evaluate the validation parameters of the modified-Agüero method, statistically significant differences (*p* < 0,05) among sets of cycle threshold values (Cts) obtained from samples analysed with the Agüero 2008 and the modified-Agüero rRT-PCRs were assessed using the *t*-test with the Microsoft Excel software. One-way ANOVA test followed by Fisher least significant difference (LSD) *post-hoc* test was used to compare the diagnostic performance of the three rRT-PCR methods, using the XLSTAT (v.2025.1.3) Excel data analysis add-on software (Lumivero (Addinsoft), 2025).

## Results

### In-silico analysis of the novel South African AHSV strain sequence

To detect mismatches that could potentially explain the lack of sensitivity of the Agüero rRT-PCR, an *in-silico* comparison of the primers/probe of the Agüero and Guthrie methods with the sequence of the undetected South African virus strain provided by the ARC-OVI was performed.

As shown in figure 1A, mismatches were detected in the reverse primer (1 position, T to C at the 14^th^ nucleotide from the 5’ primer end) and probe (2 positions, A to G and C to T at the 7^th^ and 10^th^ nucleotide from the 5’ probe end respectively) of the Agüero 2008 method. As for the Guthrie primers and probe, two mismatches were found in the forward primer, one in the probe and one in the reverse primer (figure 1B). Accordingly, modifications were made to generate the optimized reverse primer and probe based on the Agüero 2008 method (figure 1C). Subsequently, the primer/probe sequences of the original and modified methods were compared *in-silico* with 227 AHSV sequences deposited in GenBank. There were 110 and 13 AHSV sequences showing at least one mismatch with Agüero 2008 and Guthrie 2013, classified in 14 and 3 different patterns, respectively (table 1).

### Validation of the modified Agüero rRT-PCR

To assess exclusivity, other orbiviruses, several viruses affecting horses, and the most common cell lines used to propagate AHSV (sup. Table 1) were tested. Similarly to the exclusivity shown by the Agüero 2008 method, these samples were not detected with the modified-Agüero rRT-PCR (data not shown).

Analytical sensitivity was assessed by testing ten-fold serial dilutions of reference strains from all nine AHSV serotypes using both modified and original methods in parallel, in two technical replicates. All reference strains of the nine viral serotypes were properly detected by the modified method, in the same concentration range as the original method (table 2). These results show that the modified rRT-PCR exhibits a similar analytical sensitivity as compared to the original method.

**Table 2.** Analytical sensitivity of the modified-Agüero rRT-PCR compared with the Agüero 2008 method, using ten-fold serial dilutions of the nine AHSV reference strains (serotypes 1 to 9).

| DILUTION<br>(TCID<sub>50</sub>/ml) | Agüero 2008<br>(Ct) | Modified-Agüero (Ct) | DILUTION<br>(TCID<sub>50</sub>/ml) | Agüero 2008<br>(Ct) | Modified-Agüero (Ct) |
| --- | --- | --- | --- | --- | --- |
| **AHSV Serotype 1** |  |  | **AHSV Serotype 2** |  |  |
| -2 (10<sup>4.6</sup>) | 23.13 / 22.75 | 20.17 / 19.06 | -2 (10<sup>4.5</sup>) | 22.73 / 23.65 | 20.43 / 21.41 |
| -3 (10<sup>3.6</sup>) | 26.53 / 26.91 | 23.13 / 23.73 | -3 (10<sup>3.5</sup>) | 26.81 / 26.36 | 25.17 / 25.36 |
| -4 (10<sup>2.6</sup>) | 29.96 / 30.06 | 28.48 / 29.30 | -4 (10<sup>2.5</sup>) | 30.50 / 30.58 | 27.97 / 28.91 |
| -5 (10<sup>1.6</sup>) | 34.53 / 34.12 | 34.77 / 31.12 | -5 (10<sup>1.5</sup>) | 33.40 / 34.95 | 32.17 / 32.98 |
| -6 (10<sup>0.6</sup>) | 35.53 / neg | 37.62 / 37.74 | -6 (10<sup>0.5</sup>) | neg / 37.02 | 35.82 / 36.93 |
| -7 (10<sup>0.06</sup>) | neg / neg | 37.10 / neg | -7 (10<sup>0.05</sup>) | neg / neg | 36.92 / neg |
| -8 (10<sup>0.06</sup>) | neg / neg | neg / neg | -8 (10<sup>0.005</sup>) | neg / neg | neg / neg |
| **AHSV Serotype 3** |  |  | **AHSV Serotype 4** |  |  |
| -2 (10<sup>3.6</sup>) | 24.49 / 23.97 | 22.86 / 21.96 | -2 (10<sup>5.2</sup>) | 24.10 / 23.44 | 22.21 / 21.64 |
| -3 (10<sup>2.6</sup>) | 27.81 / 26.82 | 26.17 / 26.19 | -3 (10<sup>4.2</sup>) | 26.80 / 26.86 | 25.51 / 25.15 |
| -4 (10<sup>1.6</sup>) | 30.59 / 30.48 | 30.10 / 30.02 | -4 (10<sup>3.2</sup>) | 30.85 / 30.51 | 29.74 / 28.61 |
| -5 (10<sup>0.6</sup>) | 34.64 / 32.25 | 32.89 / 33.14 | -5 (10<sup>2.2</sup>) | 33.84 / 34.11 | 32.38 / 32.97 |
| -6 (10<sup>0.06</sup>) | neg / neg | 37.53 / 36.32 | -6 (10<sup>1.2</sup>) | 36.22 / neg | 36.38 / 36.95 |
| -7 (10<sup>0.006</sup>) | neg / neg | neg / neg | -7 (10<sup>0.2</sup>) | neg / neg | neg / neg |
| -8 (10<sup>0.0006</sup>) | neg / neg | neg / neg | -8 (10<sup>0.02</sup>) | neg / neg | neg / neg |
| **AHSV Serotype 7** |  |  | **AHSV Serotype 8** |  |  |
| -2 (10<sup>5.1</sup>) | 24.80 / 25.39 | 26.07 / 24.79 | -2 (10<sup>4.6</sup>) | 25.04 / 24.86 | 23.90 / 23.75 |
| -3 (10<sup>4.1</sup>) | 28.25 / 28.23 | 28.16 / 28.09 | -3 (10<sup>3.6</sup>) | 28.30 / 27.87 | 27.38 / 27.02 |
| -4 (10<sup>3.1</sup>) | 31.14 / 31.84 | 31.72 / 30.79 | -4 (10<sup>2.6</sup>) | 33.56 / 32.74 | 32.18 / 31.92 |
| -5 (10<sup>2.1</sup>) | 36.89 / 36.39 | 36.48 / 35.09 | -5 (10<sup>1.6</sup>) | 35.54 / 36.13 | 34.01 / 35.27 |
| -6 (10<sup>1.1</sup>) | neg / neg | neg / 38.62 | -6 (10<sup>0.6</sup>) | neg / neg | 38.32 / neg |
| -7 (10<sup>0.1</sup>) | neg / neg | neg / neg | -7 (10<sup>0.06</sup>) | neg / neg | neg / neg |
| -8 (10<sup>0.01</sup>) | neg / neg | neg / neg | -8 (10<sup>0.006</sup>) | neg / neg | neg / neg |
| **AHSV Serotype 5** |  |  | **AHSV Serotype 6** |  |  |
| -2 (10<sup>3.9</sup>) | 23.03 / 23.46 | 18.49 / 20.94 | -2 (10<sup>5.1</sup>) | 23.69 / 23.38 | 21.43 / 20.59 |
| -3 (10<sup>2.9</sup>) | 25.80 / 26.10 | 27.61 / 24.09 | -3 (10<sup>4.1</sup>) | 26.46 / 26.57 | 25.52 / 24.50 |
| -4 (10<sup>1.9</sup>) | 30.73 / 29.33 | 28.07 / 28.04 | -4 (10<sup>3.1</sup>) | 30.18 / 30.62 | 30.13 / 29.43 |
| -5 (10<sup>0.9</sup>) | 34.10 / 35.19 | 32.56 / 32.93 | -5 (10<sup>2.1</sup>) | 34.01 / 34.21 | 33.13 / 33.40 |
| -6 (10<sup>0.09</sup>) | neg / 37.46 | 37.32 / 37.02 | -6 (10<sup>1.1</sup>) | 37.26 / 35.97 | 36.95 / 36.25 |
| -7 (10<sup>0.009</sup>) | neg / neg | neg / neg | -7 (10<sup>0.1</sup>) | 37.41 / neg | neg / neg |
| -8 (10<sup>0.0009</sup>) | neg / neg | neg / neg | -8 (10<sup>0.01</sup>) | neg / neg | neg / neg |
| **AHSV Serotype 9** |  |  |  |  |  |
| -2 (10<sup>5.1</sup>) | 24.95 / 24.87 | 22.51 / 23.51 | TCID<sub>50</sub>, tissue culture infectious dose 50%; AHSV,<br>African Horse Sickness Virus; Ct, cycle threshold;<br>neg, negative sample. Two replicates per dilution<br>are shown. |  |  |
| -3 (10<sup>4.1</sup>) | 28.03 / 27.06 | 26.35 / 25.67 |  |  |  |
| -4 (10<sup>3.1</sup>) | 30.72 / 31.68 | 29.77 / 31.30 |  |  |  |
| -5 (10<sup>2.1</sup>) | 34.58 / 35.77 | 32.07 / 34.89 |  |  |  |
| -6 (10<sup>1.1</sup>) | neg / neg | 38.47 / 37.78 |  |  |  |
| -7 (10<sup>0.1</sup>) | neg / neg | neg / neg |  |  |  |
| -8 (10<sup>0.01</sup>) | neg / neg | neg / neg |  |  |  |

Inclusivity was evaluated by analysing viral suspensions from the AHSV EURL’s strain collection, showing that all the strains were properly detected by the modified-Agüero method and similar Ct values were obtained with both methods (Mean Ct = 21.33 ± 4.79 SD vs. 21.89 ± 5.31 SD; *p*-value= 0.64; sup. Table 2).

To further ensure the inclusivity of the modified-Agüero method, additional strains of the AHSV collection maintained at Pirbright were analysed. To this end, sequences of the modified-Agüero primers/probe were provided to the WOAH Reference laboratory for AHS in the UK, where analyses of AHSV strains of their virus collection showing mismatches with primers/probes of the reference methods were carried out. The sequences of these AHSV strains correspond to some strain patterns described in table 1 and some strains that have additional mismatches (table 3).

**Table 3.** Comparative assessment of the Agüero, modified-Agüero and Guthrie 2013 methods using the TPI AHSV collection strains.

| Virus strain | Agüero 2008 |  | Modified-Agüero |  | Guthrie 2013 |  |
| --- | --- | --- | --- | --- | --- | --- |
|  | Mismatches* | Ct | Mismatches* | Ct | Mismatches* | Ct |
| ETH 2010/18 | Pattern A1 | 22.54 | Pattern A1 | 22.45 | No mismatches | Nd |
| SEN 1998/01 | Pattern A3 plus R 7 (A>G) | 35.33 | Pattern A3 plus R 7 (A>G) | 35.05 | R 21 (A>G) | 31.86 |
| ETH 2010/08 | Pattern A4 plus R 4 (A>G) | 27.20 | Pattern A4 plus R 4 (A>G) | 27.07 | Pattern G2 | 22.49 |
| THA 2020/01 | Pattern A5 | 24.86 | Pattern A5 | 25.11 | No mismatches | Nd |
| ETH 2010/13 | Pattern A11 | 31.19 | No mismatches | 30.24 | Pattern G1 | 28.53 |
| ETH 2019/02 |  | 25.21 |  | 24.24 |  | 23.47 |
| ETH 2019/03 | Pattern A11 plus R 14 (T>C) | 24.89 | No mismatches | 23.71 | Pattern G1 plus P 10 (A>G) | 22.88 |
| ETH 2019/07 |  | 27.84 |  | 26.71 |  | 25.80 |
| ETH 2019/01 |  | Nd |  | 23.35 |  | 20.61 |
| ETH 2019/04 |  | Nd |  | 23.81 |  | 21.00 |
| ETH 2019/05 | No mismatches | Nd | No mismatches | 23.79 | Pattern G3 | 21.17 |
| ETH 2019/06 |  | Nd |  | 24.97 |  | 22.30 |

\* Described in relation to patterns included in table 1, the position of mismatches from 5' in primer F (F), primer R (R) or probe (P) is indicated; Ct, cycle threshold; Nd: Not done (modified-Agüero and Guthrie rRT-PCRs were not performed in strains with no mismatches); ETH Ethiopia; THA, Thailand; SEN, Senegal. Agüero 2008 and Guthrie 2013 assays were only performed on strains that had mismatches with the primers and/or probe of that specific rRT-PCR.

As shown in table 3, the three methods detected the AHSV strains presenting one or two mismatches with the primers or probe. Likewise, viral strains showing one mismatch in any of the primers/probe or one mismatch in a primer plus a mismatch in the other primer or the probe, have been properly detected with the three rRT-PCR methods.

Regarding diagnostic performance, similar Ct values were obtained with both methods from convalescent horses’ EDTA blood (Agüero 2008 mean Ct = 28.31 ± 2.34 SD vs. modified-Agüero mean Ct = 27.64 ± 2.44 SD; *p*-value= 0.31) and tissue samples (Agüero 2008 mean Ct = 26.18 ± 4.2 SD vs. modified-Agüero mean Ct = 25.99 ± 4.3 SD; *p*-value= 0.88) obtained during the Kenya outbreaks (sup. Table 3). In addition, twenty-four (24) spleen samples from affected horses in Spain (1989-90 outbreaks) were analysed using both methods (sup. Table 4). Again, comparable Ct values were obtained (Agüero 2008 mean Ct = 27.34 ± 4.15 SD vs. modified-Agüero method mean Ct = 26.37 ± 3.41 SD; *p*- value = 0.38). Thus, our data demonstrate that the diagnostic sensitivity of the modified-Agüero rRT-PCR is comparable to that of the Agüero 2008 method in clinical samples (blood and tissues). Overall, from seventy-five (75) analysed positive clinical samples, the agreement in the results was 100% without significant differences in the Ct values obtained.

**Table 4.**
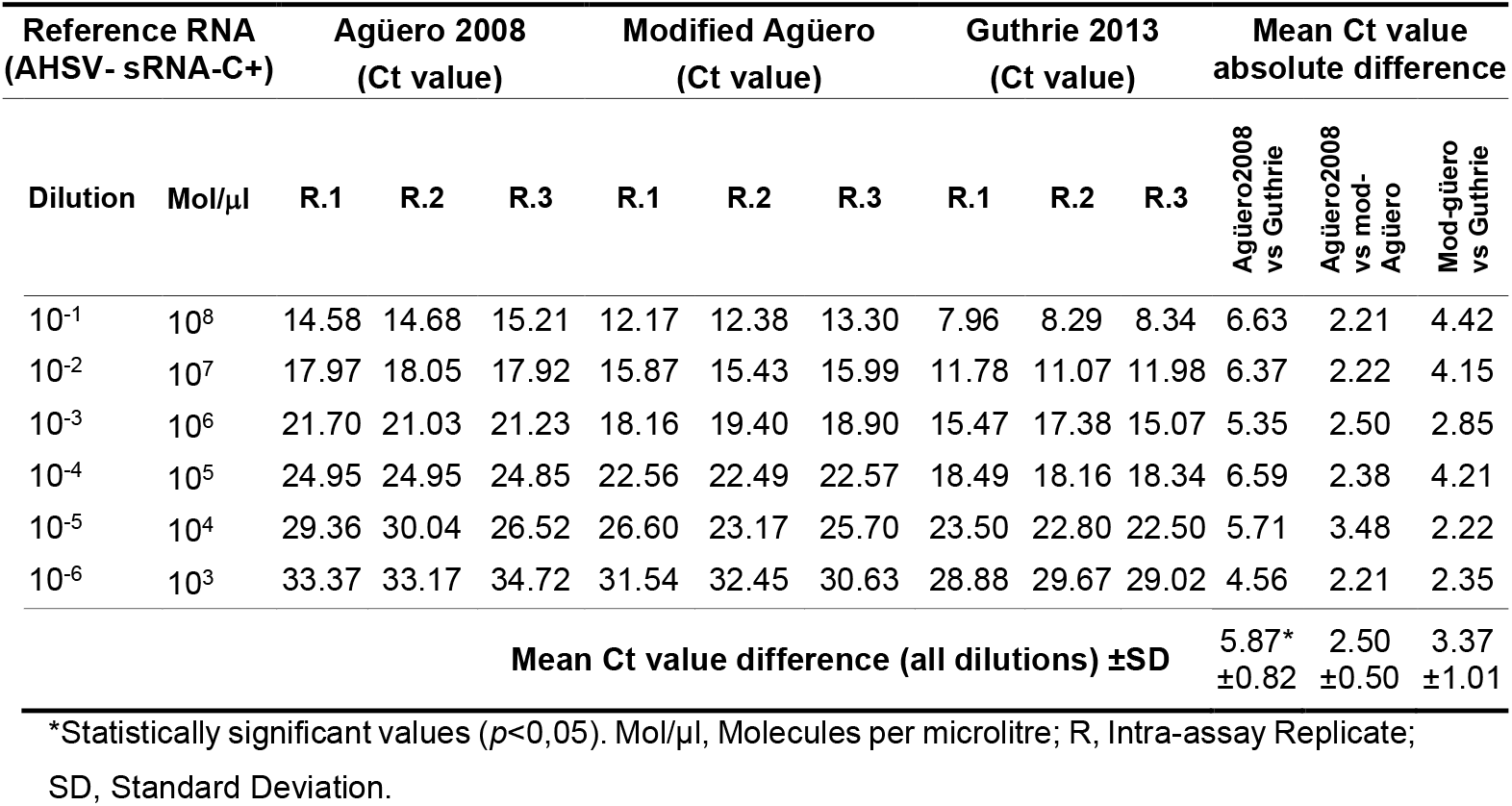
Comparative assessment of the Agüero 2008, modified-Agüero and Guthrie 2013 rRT-PCRs using the synthetic RNA containing the segment-7 target sequence of the AHSV reference strain (AHSV-sRNA-C+).

To check diagnostic specificity, twenty-eight (28) EDTA blood samples from horses from an AHS-free country were analysed. As expected, both methods yielded negative results (data not shown).

To complete the evaluation of the modified-Agüero rRT-PCR validation parameters, intra-assay repeatability was assessed. The observed variation in Ct values between technical replicates was low (mean absolute difference in Ct value = 0.9 ± 0.84 SD) (sup. Table 5).

**Table 5.**
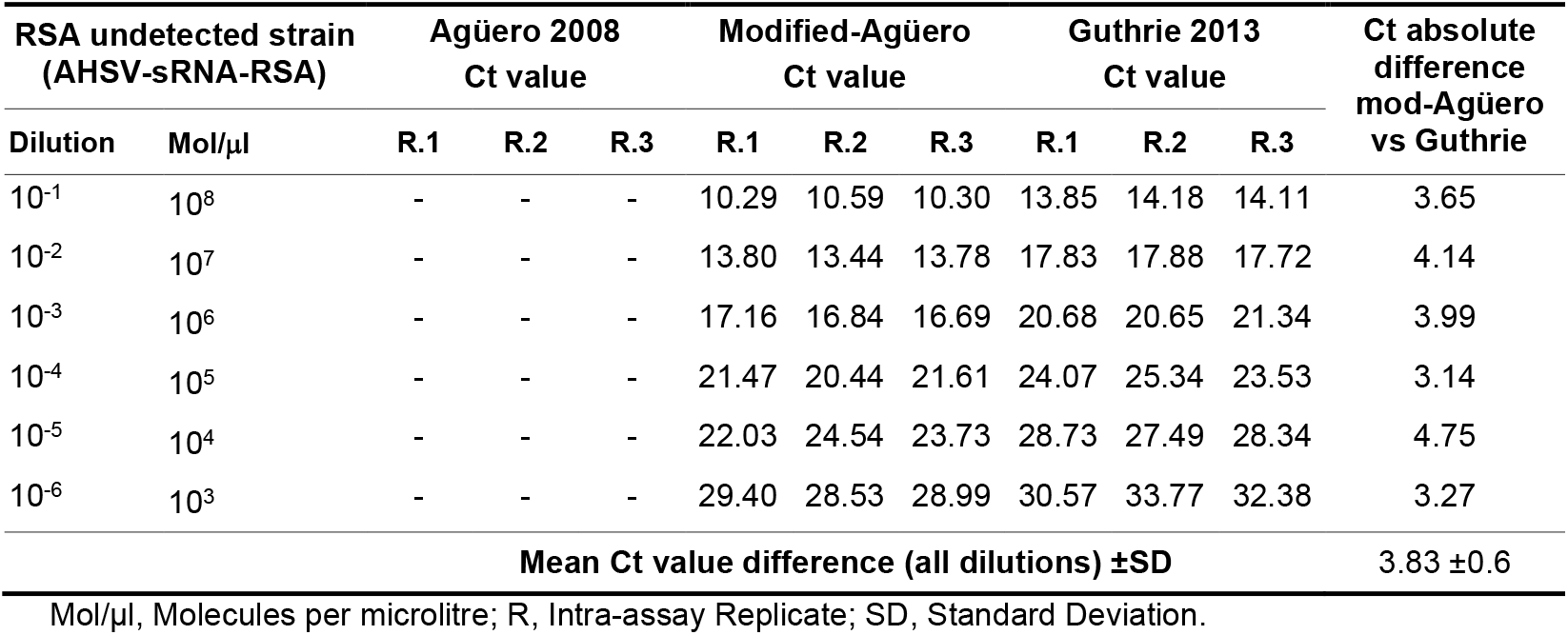
Comparative assessment of the Agüero 2008, modified-Agüero and Guthrie 2013 rRT-PCRs using the synthetic RNA containing the segment-7 target sequence of the South African undetected strain (AHSV-sRNA-RSA).

| RSA undetected strain<br>(AHSV-sRNA-RSA) |  | Agüero 2008<br>Ct value |  |  | Modified-Agüero<br>Ct value |  |  | Guthrie 2013<br>Ct value |  |  | Ct absolute<br>difference<br>mod-Agüero<br>vs Guthrie |
| --- | --- | --- | --- | --- | --- | --- | --- | --- | --- | --- | --- |
| Dilution | Mol/μl | R.1 | R.2 | R.3 | R.1 | R.2 | R.3 | R.1 | R.2 | R.3 |  |
| 10<sup>-1</sup> | 10<sup>8</sup> | - | - | - | 10.29 | 10.59 | 10.30 | 13.85 | 14.18 | 14.11 | 3.65 |
| 10<sup>-2</sup> | 10<sup>7</sup> | - | - | - | 13.80 | 13.44 | 13.78 | 17.83 | 17.88 | 17.72 | 4.14 |
| 10<sup>-3</sup> | 10<sup>6</sup> | - | - | - | 17.16 | 16.84 | 16.69 | 20.68 | 20.65 | 21.34 | 3.99 |
| 10<sup>-4</sup> | 10<sup>5</sup> | - | - | - | 21.47 | 20.44 | 21.61 | 24.07 | 25.34 | 23.53 | 3.14 |
| 10<sup>-5</sup> | 10<sup>4</sup> | - | - | - | 22.03 | 24.54 | 23.73 | 28.73 | 27.49 | 28.34 | 4.75 |
| 10<sup>-6</sup> | 10<sup>3</sup> | - | - | - | 29.40 | 28.53 | 28.99 | 30.57 | 33.77 | 32.38 | 3.27 |
| Mean Ct value difference (all dilutions) ±SD |  |  |  |  |  |  |  |  |  |  | 3.83 ±0.6 |

Mol/μl, Molecules per microlitre; R, Intra-assay Replicate; SD, Standard Deviation.

### Comparative assessment to detect the new AHSV variant from RSA

To test the ability of the modified-Agüero method to detect the new AHSV variant from RSA, a comparative assessment with the Agüero 2008 and the Guthrie 2013 method was performed using the synthetic RNAs mimicking the AHSV reference strain (AHSV-sRNA-C+) and the RSA undetected AHSV strain (AHSV-sRNA-RSA) as target sequences. The synthetic RNAs were used to prepare a solution containing 10^9^ molecules/μl in water, then, ten-fold serial dilutions were obtained and 2 μl of each dilution was tested with the three rRT-PCR methods. As expected, the three methods detected the positive control (AHSV-sRNA-C+) (table 4). Specifically, the Guthrie 2013 method presented a statistically significant lower mean Ct value to detect this AHSV-sRNA-C+ as compared to the Agüero 2008 rRT-PCR (mean Ct value difference = 5.87* ± 0.82 SD, *p*-value = 0.012). No statistically significant differences were observed between the Ct values obtained with the modified-Agüero method and the WOAH reference rRT-PCRs (mean Ct value difference with the Guthrie method = 3.37 ± 1.01 SD, *p*-value = 0.141; mean Ct value difference versus the Agüero 2008 method = 2.50 ± 0.50 SD, *p*-value =0.272).

As shown in table 5, the AHSV-sRNA-RSA was undetected by the Agüero 2008 rRT-PCR whereas it was positively ascertained by the modified-Agüero and the Guthrie 2013 methods. Interestingly, the observed Ct values using the modified-Agüero method were consistently lower than those obtained by the Guthrie rRT-PCR, independently of the target AHSV-sRNA-RSA dilution (mean Ct value difference = 3.83 ± 0.6 SD *p*-value =0.081).

## Discussion

Given the relevance of AHS, effective vigilance, rapid response, and collaboration to reduce its impact and protect equine populations are key to minimize the risks posed by the disease to animal health and the equine industry worldwide (Castillo-Olivares, 2021; Dennis et al., 2019; Redmond et al., 2022). To meet these goals, the availability of validated and harmonised AHS diagnostic methods is essential. The Agüero 2008 is one of the two rRT-PCR methods for AHSV molecular diagnosis included in the WOAH manual (World Organisation for Animal Health, 2025a), and it is used worldwide. Moreover, the EU Regulation 2016/429 of the European Parliament and of the Council, article 94 point 1, states that EURLs shall contribute to the improvement and harmonization of methods of analysis, test or diagnosis to be used by official laboratories designated (European Union, 2016). Therefore, upon notification by the WOAH reference laboratory in RSA that the Agüero 2008 rRT-PCR was unable to detect a new AHSV strain from Western Cape, as EURL and WOAH reference laboratory for AHS, we conducted the modification and validation of the Agüero 2008 rRT-PCR method.

The *in-silico* comparison of the primers/probe of the Agüero 2008 and Guthrie 2013 rRT-PCRs with the sequence of the undetected South African virus provided by the RSA WOAH reference laboratory revealed mismatches with the reverse primer (1 position) and probe (2 positions) of the Agüero 2008 method. Since PCR with degenerate primers can be used to detect genetic variants (Campos & Quesada, 2017), we optimized the sensitivity of the Agüero 2008 method by introducing degenerate nucleotides at the mismatch positions of the reverse primer and probe. These changes allowed a better complementarity and annealing than the Agüero 2008 reverse primer and probe with the novel RSA AHSV strain segment-7 target sequence, while maintaining a good complementarity with the target sequence of AHSV reference and collection strains.

We demonstrated that the modified assay exhibited a similar analytical sensitivity as the original Agüero 2008 method, while analytical specificity (exclusivity) was maintained. Inclusivity was evaluated by using an extensive AHSV reference collection from the EURL and The Pirbright Institute, including AHSV strains showing some mismatches in primers and or probe. Moreover, the diagnostic performance results obtained using samples from convalescent horses taken during AHS outbreaks and from negative horses from AHS-free zones, showed that the modified-Agüero method presents an equivalent diagnostic specificity and sensitivity as compared to the original Agüero 2008 rRT-PCR. Overall, the data obtained from the comparative analyses demonstrated that the modified-Agüero 2008 rRT-PCR shows similar analytical and diagnostic performance, as compared with the original Agüero 2008 method. Also, the observed variation in Ct values between replicates was low, showing a good repeatability of the assay.

Although the RSA WOAH reference laboratory shared the sequence of the new AHSV RSA strain targeted by the Agüero 2008 and Guthrie 2013 rRT-PCRs, the shipment of this virus to the AHS EURL was not possible. Since synthetic RNAs have been successfully used as spike-in controls in RNA-seq experiments (Jiang et al., 2011), we designed a synthetic RNA of 120 nucleotides containing the amplicon generated by the Agüero 2008 method (AHSV-sRNA-RSA). This synthetic nucleic acid was used to carry out a comparative assessment of the diagnostic performance of the modified-Aguero method versus the WOAH reference rRT-PCR methods.

The results of the comparative assessment of the modified-Agüero versus the WOAH manual reference methods showed that all three rRT-PCRs detected the positive control AHSV-sRNA-C+. In this regard, our data shows a lower Ct value of the Guthrie 2013 method to this positive control as compared to the Agüero 2008 rRT-PCR, and a similar sensitivity to the modified-Agüero rRT-PCR. With respect to the AHSV-sRNA-RSA, it was not detected by the Agüero 2008 rRT-PCR but positively identified by the modified-Agüero and the Guthrie 2013 methods. The explanation on why the Agüero 2008 original method with a total of three mismatches (one in the reverse primer and two in the probe) was unable to detect the relevant RSA strain, whereas the Guthrie method with five mismatches (two in the forward primer, one in the probe and two in the reverse primer) was able to detect the RSA strain remains hitherto elusive. A plausible explanation is that the effects of mismatches in the probe binding region could have a more pronounced effect for a broad range of characteristics of real-time PCR amplification curves than mismatches in the primer (Süss et al., 2009). Therefore, the original Agüero 2008 method, with two mismatches in the probe, could be more impaired to detect the new RSA AHSV strain than the Guthrie method. Interestingly, the observed Ct values obtained using the modified-Agüero method were lower than those attained with the Guthrie rRT-PCR at any target AHSV-sRNA-RSA dilution, suggesting a higher sensitivity of the modified-Agüero versus the Guthrie method to detect this strain. This can be due to the better complementarity of the optimized primer and probe of the modified-Agüero method with the new RSA AHSV strain target sequence.

To further ensure that the modified-Agüero method was able to detect a wide range of reference AHSV strains, particularly those including mismatches with the primers and/or probes of any of the three methods, the diagnostic performance of the modified-Agüero rRT-PCR was evaluated and compared with that of the WOAH reference rRT-PCRs using several strains of AHSV collection from Pirbright’s Orbivirus reference collection. These assays were part of the inclusivity assessment of the modified-Agüero method and were only performed on virus strains that had mismatches with the primers and/or probe of that specific rRT-PCR, which included strains from: Senegal (Mertens & Attoui, n.d.), AHS outbreaks in Ethiopia 2010 (Aklilu et al., 2014) and Thailand 2020 (King et al., 2020). These data show that the three methods detect the AHSV strains from the Pirbright Orbivirus reference collection despite the presence of one or two mismatches in the primer and/or probe and further reinforces the adequate diagnostic performance of the modified-Agüero method.

It is noteworthy that, even though segment 7 is highly conserved in AHSV, almost half of the AHSV strains deposited in GenBank from 2009 and 2023 have at least one mismatch with respect to the primers and/or probe sequences of the Agüero method, and 13 AHSV strains have mismatches with respect to the primers and/or probe of the Guthrie method. However, no detection failures with the reference methods have been reported to date. The new AHSV strain from RSA not detected by the Agüero 2008 method is the only one that presents two mismatches in the probe, which, in agreement with Süss and cols., suggests that this might be critical for the rRT-PCR’s sensitivity.

The two reference methods have proven to be the most sensitive and specific of all published methods, as described in the WOAH Manual (World Organisation for Animal Health, 2025a). However, both methods target the same region of segment 7, and even the amplicons generated by both methods have overlapping regions. Considering that the occurrence of AHSV strains with mutations in the sequence of this area seems higher than expected for such a conserved segment, it seems advisable to have available additional rRT-PCR methods with similar sensitivity targeting other segments of the AHSV genome, which could be used in case of suspected detection failure. In this regard, there are two rRT-PCR methods targeting segments 1 and 9 currently in development with promising results (Hofmann et al, personal communication).

In conclusion, this study shows that the modified-Agüero rRT-PCR, allows the reliable detection of a wide collection of AHSV strains and clinical samples and a synthetic RNA mimicking the new AHSV strain from the RSA, the only one that has been reported so far as undetected by the Agüero 2008 original method. Therefore, we will propose substituting the Agüero 2008 AHS rRT-PCR by the modified-Agüero method in the WOAH Manual. Finally, we recommend to AHS EU-NRLs and official diagnostic laboratories worldwide, the implementation of the modified-Agüero method described here. This work highlights the relevance of the collaboration between WOAH Reference Laboratories to provide effective and robust diagnostic tools, updated to identify novel viral strains.

## Notes

### Competing Interest Statement

The authors have declared no competing interest.

### Funding

- Spanish Ministry of Agriculture, Fisheries and Food (MAPA).
- European Health and Digital Executive Agency (HADEA) (‘EU executive agency’), under the powers delegated by the European Commission

