## Supplementary material for "Modification and validation of a reference real-time RT-PCR method for the detection of a new African horse sickness virus variant"

**Supplementary figure 1. Synthetic RNA sequences and features.**

**AHS SEGMENT-7 (VP7) target sequence of the RSA strain (AHSV-sRNA-RSA)**

5'-aau ggu guu guu gcg cca gua ggc cag auc aac aga gcu cuu gug cua gcg gcu uac cac uag ugg cug cgg ugu ugc acg guc gcc gcu uuc auu agu guc gcg ucg gcu cuu aug cug-3'

Base Count: 120
Purification method: HPLC
Molecular Weight (Da): 38.508
Tm(ºC): 91
$OD_{260nm}$: 1,2
nmol: 1,0
µl required for 100µM solution: 10

**AHS SEGMENT-7 (VP7) target sequence of the AHSV reference strains (AHSV-sRNA-C+)**

5'-aau ggu guu guu gcg cca gua ggc cag auc aac aga gcu cuu gug cua gca gcc uac cac uag ugg cug cgg ugu ugc acg guc acc gcu uuc auu agu guc gcg ucg guu cuu aug cug-3'

Base Count: 120
Purification method: HPLC
Molecular Weight (Da): 38.476
Tm(ºC): 90
$OD_{260nm}$: 1,2
nmol: 1,0
µl required for 100µM solution: 10

AHS, African Horse Sickness; VP7, viral protein 7; AHSV, African horse sickness virus; AHSV-RSA-sRNA, synthetic RNA containing the target sequence of the new South African strain (highlighted in red); AHSV-sRNA-C+, synthetic RNA containing the target sequence the AHSV reference strains (highlighted in red); HPLC, High performance liquid chromatography; Da, Dalton; Tm, melting temperature; OD, optic density.

### Supplementary tables.

#### Supplementary Table 1. Details of strains and cell lines included to check exclusivity

| Virus / Cell line | Strain (Source) |
|---|---|
| Bluetongue virus (All 24 notified serotypes) | TPI, UK |
| Equine Encephalosis virus (EEV3 GAM 2009/06) | TPI, UK |
| Epizootic hemorrhagic disease virus (EHDV1 15.02 [5374]) | TPI, UK |
| Equine rhinopneumonitis type 1 | LCV, Spain |
| Equine rhinopneumonitis type 4 | LCV, Spain |
| West Nile virus (Lineage 1 Egypt 101 | Institut Pasteur, France |
| Usutu virus | LCV, Spain |
| Equine Eastern Encephalomyelitis | Anses, France |
| Western Encephalomyelitis | Anses, France |
| Venezuelan encephalomyelitis | Anses, France |
| Uninfected Vero cells suspension | ATCC |
| Uninfected BHK cells suspension | ATCC |
| Uninfiected KC cells suspension | ATCC |

ATCC, American Type Culture Collection; TPI, The Pirbright Institute; LCV, Laboratorio Central de Veterinaria; ANSES, (*Agence nationale de sécurité sanitaire de l'alimentation, de l'environnement et du travail)*; BTV, Bluetongue virus; EHDV, Epizootic haemorrhagic disease virus; Neg, negative; POS, positive (Ct value)

**Supplementary Table 2. Diagnostic sensitivity of the modified-Agüero method in viral suspensions from the AHSV strain collection maintained at the LCV**

| Strain (source) | Agüero 2008 rRT-PCR | | Modified-Agüero rRT-PCR | |
|---|---|---|---|---|
| | Result | Ct | Result | Ct |
| SEN2007 st2 (TPI) | POS | 18.44 | POS | 18.53 |
| SEN2007 st7 (TPI) | POS | 18.18 | POS | 18.08 |
| KEN2006 st9 (TPI) | POS | 18.88 | POS | 19.04 |
| KEN2007 st4 (TPI) | POS | 21.00 | POS | 21.98 |
| GHA2010 st2 (TPI) | POS | 31.51 | POS | 35.32 |
| ETH2010 st6 (TPI) | POS | 19.19 | POS | 19.34 |
| ETH2010 st2 (TPI) | POS | 24.37 | POS | 23.98 |
| ETH2010 st4 (TPI) | POS | 32.68 | POS | 31.90 |
| ETH2010 st8 (TPI) | POS | 23.64 | POS | 22.08 |
| ETH2010 st9 (TPI) | POS | 24.64 | POS | 22.93 |
| KEN st1 (CVR) | POS | 21.40 | POS | 20.99 |
| KEN st4 (CVR) | POS | 19.65 | POS | 19.92 |
| KEN st5 (CVR) | POS | 16.87 | POS | 17.19 |
| KEN st7 (CVR) | POS | 17.16 | POS | 17.30 |
| KEN st8 (CVR) | POS | 20.53 | POS | 20.37 |
| KEN st9 (CVR) | POS | 26.83 | POS | 26.72 |
| KEN2015 st2 (CVR)* | POS | 18.29 | POS | 17.81 |
| KEN2013 st4 (CVR) | POS | 20.29 | POS | 20.03 |
| KEN2015 st7 (CVR)* | POS | 18.60 | POS | 18.43 |
| KEN2016 st4 (CVR)* | POS | 22.38 | POS | 23.00 |
| KEN2017 st5 (CVR)* | POS | 19.48 | POS | 19.66 |
| SPA1988 st4 (LCV) | POS | 16.25 | POS | 15.54 |
| SPA1989 st4 (LCV) | POS | 18.45 | POS | 28.11 |
| SPA1990 st4 (LCV) | POS | 21.84 | POS | 21.26 |
| AHSV1 LAV (OBP)* | POS | 14.40 | POS | 12.20 |
| AHSV2 LAV (OBP)* | POS | 17.58 | POS | 18.39 |
| AHSV3 LAV (OBP)* | POS | 17.73 | POS | 19.12 |
| AHSV4 LAV (OBP)* | POS | 31.30 | POS | 30.91 |
| AHSV6 LAV (OBP)* | POS | 26.69 | POS | 32.31 |
| AHSV7 LAV (OBP)* | POS | 18.17 | POS | 18.70 |
| AHSV8 LAV (OBP)* | POS | 17.74 | POS | 20.57 |
| THA2020 st1 (NIAH)* | POS | 31.78 | POS | 31.54 |
| SPA1988 st4 (LCV O65) | POS | 19.78 | POS | 19.67 |
| SPA1987 st4 (LCV O66) | POS | 19.49 | POS | 21.46 |
| | **Mean Ct** | **21.33** | **Mean Ct** | **21,89** |
| | **SD** | **4.79** | **SD** | **5,31** |
| | | | **Mean Ct absolute difference** | **0,56** |
| | | | ***p*-value** | **0.142** |

AHSV, African Horse Sicknes Virus; st, serotype; POS, positive sample; Ct, cycle threshold; SD, standard deviation; SEN, Senegal; GHA, Ghana; KEN, Kenya; SPA, Spain; R., Sample registry number at the LCV; (OBP), Onderstepoort Biological Products soc ltd. TPI: The Pirbright Institute; CVR: Central Veterinary Research Laboratory; LCV: Laboratorio Central de Veterinaria; NIAH: National Institute of Animal Health (Thailand); B: BHK; V: Vero; *Virus isolated in the LCV from clinical samples received from this source.

**Supplementary Table 3. Diagnostic performance of the modified-Agüero method in EDTA-blood and tissue samples from convalescent horses obtained during the Kenya outbreak**

| Identification | Type | Serotype | Agüero 2008 | | Modified-Agüero | |
|---|---|---|---|---|---|---|
| | | | Result | Ct | Result | Ct |
| 2441/15 17 | B | 4 | POS | 30,68 | POS | 28,34 |
| 2441/15 19 | B | 2 | POS | 26,92 | POS | 26,2 |
| 2441/15 20 | B | 5 | POS | 31,37 | POS | 31,13 |
| 2441/15 13 | B | 5 | POS | 24,89 | POS | 23,48 |
| 2782/15 1 | B | 2 | POS | 29,89 | POS | 28,95 |
| 2782/15 2 | B | 2 | POS | 32,52 | POS | 33,17 |
| 2782/15 3 | B | 2 | POS | 24,97 | POS | 25,75 |
| 56/16 2 | B | 4 | POS | 26,97 | POS | 25,66 |
| 256/16 3 | B | 9 | POS | 26,88 | POS | 26,85 |
| 256/16 4 | B | 7 | POS | 27,26 | POS | 26,04 |
| 377/16 2 | B | 4 | POS | 29,05 | POS | 28,88 |
| 377/16 3 | B | 4 | POS | 26,39 | POS | 25,52 |
| 841/16 8 | B | 4 | POS | 28,04 | POS | 27,14 |
| 841/16 9 | B | 7 | POS | 25,26 | POS | 24,52 |
| 841/16 10 | B | 2 | POS | 28,1 | POS | 28,09 |
| 1766/16 2 | B | 2 | POS | 29,4 | POS | 28,31 |
| 1766/16 3 | B | 2 | POS | 29,15 | POS | 28,26 |
| 1766/16 4 | B | 2 | POS | 29,12 | POS | 27,94 |
| 1766/16 5 | B | 4 | POS | 29,52 | POS | 28,44 |
| 2459/16 2 | B | 4 | POS | 32,29 | POS | 30,74 |
| 2459/16 3 | B | 9 | POS | 30,03 | POS | 27,59 |
| 2459/16 4 | B | 4 | POS | 28,09 | POS | 26,95 |
| 2459/16 5 | B | 3 | POS | 30,65 | POS | 30,25 |
| 2772/16 1 | B | 9 | POS | 25,19 | POS | 22,34 |
| 1083/17 1 | B | 5 | POS | 23,95 | POS | 27,2 |
| 1838/17 1 | B | 5 | POS | 29,57 | POS | 30,87 |
| 2441/15 (11) | S | 5 | POS | 25,39 | POS | 27,52 |
| 2441/15 (16) | S | 4 | POS | 28,71 | POS | 28,34 |
| 2782/15 (4) | Li | 2 | POS | 28,95 | POS | 28,22 |
| 2782/15 (5) | S | 2 | POS | 27,33 | POS | 26,2 |
| 2782/15 (6) | Lu | 2 | POS | 20 | POS | 18,65 |
| 256/16 (5) | Lu | 4 | POS | 33,38 | POS | 31,98 |
| 256/16 (6) | H | 4 | POS | 32,54 | POS | 21,8 |
| 256/16 (7) | S | 4 | POS | 24,7 | POS | 24,08 |
| 256/16 (8) | Li | 4 | POS | 25,66 | POS | 23,77 |
| 256/16 (9) | Lu | 9 | POS | 31,56 | POS | 30,77 |
| 256/16 (10) | H | 9 | POS | 22,4 | POS | 20,88 |
| 256/16 (11) | S | 9 | POS | 27,16 | POS | 24,78 |
| 256/16 (12) | Li | 9 | POS | 26,35 | POS | 23,8 |
| 1766/16 (8) | H | 4 | POS | 29,79 | POS | 28,94 |
| 1766/16 (9) | Lu | 4 | POS | 33,67 | POS | 32,61 |
| 2459/16 (6) | Lu | 7 | POS | 24,49 | POS | 22,7 |
| 2459/16 (9) | Li | 7 | POS | 26,48 | POS | 24,49 |
| 2772/16 (3) | H | 4 | POS | 26,85 | POS | 26,29 |
| 2772/16 (4) | S | 4 | POS | 26,95 | POS | 33,95 |
| 2772/16 (5) | S | 4 | POS | 26,53 | POS | 30,81 |
| 1083/17 (3) | Lu | 5 | POS | 21,53 | POS | 23,65 |
| 1083/17 (5) | S | 5 | POS | 23,76 | POS | 26,26 |
| 1083/17 (7) | Lu | 5 | POS | 17,46 | POS | 18,52 |
| 1083/17 (8) | H | 5 | POS | 23,21 | POS | 30,15 |
| 1083/17 (9) | Li | 5 | POS | 19,55 | POS | 20,7 |

| | | | | |
|---|---|---|---|---|
| Statistical analysis of blood samples | Mean Ct | 28,31 | Mean Ct | 27,64 |
| | SD | 2,34 | SD | 2,44 |
| | Mean Ct absolute difference | | | 0,67 |
| | | | *p*-value | 0,31 |
| Statistical analysis of tissue samples | Mean | 26.18 | Mean | 25.99 |
| | SD | 4.20 | SD | 4.3 |
| | Mean Ct absolute difference | | | 0.18 |
| | | | *p*-value | 0,88 |

**POS, positive sample; Ct, cycle threshold; SD, standard deviation; B, blood-EDTA; S, spleen; Li, liver; Lu, Lung; H, heart.**

**Supplementary Table 4. Diagnostic performance of the modified Agüero method in blood EDTA samples from affected horses in Spain during the (1989-90) outbreak**

| Identification | Agüero 2008 | | Agüero-New-mod | |
|---|---|---|---|---|
| | **Result** | **Ct** | **Result** | **Ct** |
| 1542/89 (44) | POS | 20,3 | POS | 20,42 |
| 1542/89 (44) | POS | 22,1 | POS | 22,72 |
| 1542/89 (43) | POS | 20,4 | POS | 21,57 |
| 1542/89 (37) | POS | 22,7 | POS | 23,62 |
| 1542/89 (33) | POS | 22,9 | POS | 20,96 |
| 1542/89 (32) | POS | 23,2 | POS | 22,91 |
| 1542/89 (28) | POS | 24,8 | POS | 24,53 |
| 1542/90 (41) | POS | 25,1 | POS | 24,94 |
| 1371/89 ( 51) | POS | 31,1 | POS | 29,17 |
| 1371/89 ( 51) | POS | 28,8 | POS | 28,01 |
| 1371/89 ( 5) | POS | 30,2 | POS | 29,37 |
| 1371/89 ( 55) | POS | 24,1 | POS | 24,06 |
| 1371/89 ( 6) | POS | 30,6 | POS | 27,84 |
| 1371/89 ( 67) | POS | 29,3 | POS | 27,57 |
| 1371/89 ( 8) | POS | 28,5 | POS | 26,53 |
| 1371/89 ( 35) | POS | 32,2 | POS | 31,13 |
| 1371/89 ( 28) | POS | 33,7 | POS | 32,28 |
| 1371/89 ( 3) | POS | 31,9 | POS | 30,88 |
| 1378/89 ( 41) | POS | 23,6 | POS | 23,07 |
| 1378/89 ( 7) | POS | 29,5 | POS | 29,61 |
| 1378/89 ( 71) | POS | 31,1 | POS | 27,58 |
| 1378/89 ( 46) | POS | 27,3 | POS | 25,61 |
| 1378/89 ( 32) | POS | 31,4 | POS | 30,28 |
| 1378/89 ( 15) | POS | 31,3 | POS | 28,23 |
| | **Mean Ct** | 27,34 | **Mean Ct** | 26,37 |
| | **SD** | 4,15 | **SD** | 3,46 |
| | **Mean Ct absolute difference** | | | 0,97 |
| | ***p*-value** | | | 0,38 |

POS, positive sample; Ct, cycle threshold; SD, standard deviation

### Supplementary table 5. Intra-assay repeatability of the modified-Agüero rRT-PCR method

| AHSV prototype strain | Dilution | Rep1 | Rep2 | Absolute Ct value difference | AHSV prototype strain | Dilution | Rep1 | Rep2 | Absolute Ct value difference |
|---|---|---|---|---|---|---|---|---|---|
| AHSV-1 | -2($10^{4.6}$) | 20,17 | 19,06 | 1,11 | AHSV-2 | -2($10^{4.5}$) | 20,43 | 21,41 | 0,98 |
| | -3 ($10^{3.6}$) | 23,13 | 23,73 | 0,6 | | -3 ($10^{3.5}$) | 25,17 | 25,36 | 0,19 |
| | -4 ($10^{2.6}$) | 28,48 | 29,3 | 0,82 | | -4 ($10^{2.5}$) | 27,97 | 28,91 | 0,94 |
| | -5 ($10^{1.6}$) | 34,77 | 31,12 | 3,65 | | -5 ($10^{1.5}$) | 32,17 | 32,98 | 0,81 |
| | -6 ($10^{0.6}$) | 37,62 | 37,74 | 0,12 | | -6 ($10^{0.5}$) | 35,82 | 36,93 | 1,11 |
| | -7 ($10^{0.06}$) | 37,1 | neg | | | -7($10^{0.05}$) | 36,92 | neg | |
| | -8 ($10^{0.06}$) | neg | neg | | | -8($10^{0.005}$) | neg | neg | |
| AHSV-3 | -2($10^{3.6}$) | 22,86 | 21,96 | 0,9 | AHSV-4 | -2($10^{5.2}$) | 22,21 | 21,64 | 0,57 |
| | -3 ($10^{2.6}$) | 26,17 | 26,19 | 0,02 | | -3 ($10^{4.2}$) | 25,51 | 25,15 | 0,36 |
| | -4 ($10^{1.6}$) | 30,1 | 30,02 | 0,08 | | -4 ($10^{3.2}$) | 29,74 | 28,61 | 1,13 |
| | -5 ($10^{0.6}$) | 32,89 | 33,14 | 0,25 | | -5 ($10^{2.2}$) | 32,38 | 32,97 | 0,59 |
| | -6 ($10^{0.06}$) | 37,53 | 36,32 | 1,21 | | -6 ($10^{1.2}$) | 36,38 | 36,95 | 0,57 |
| | -7 ($10^{0.006}$) | neg | neg | | | -7 ($10^{0.2}$) | neg | neg | |
| | -8($10^{0.0006}$) | neg | neg | | | -8 ($10^{0.02}$) | neg | neg | |
| AHSV-5 | -2($10^{3.9}$) | 18,49 | 20,94 | 2,45 | AHSV-6 | -2($10^{5.1}$) | 21,43 | 20,59 | 0,84 |
| | -3 ($10^{2.9}$) | 27,61 | 24,09 | 3,52 | | -3 ($10^{4.1}$) | 25,52 | 24,5 | 1,02 |
| | -4 ($10^{1.9}$) | 28,07 | 28,04 | 0,03 | | -4 ($10^{3.1}$) | 30,13 | 29,43 | 0,7 |
| | -5 ($10^{0.9}$) | 32,56 | 32,93 | 0,37 | | -5 ($10^{2.1}$) | 33,13 | 33,4 | 0,27 |
| | -6 ($10^{0.09}$) | 37,32 | 37,02 | 0,3 | | -6 ($10^{1.1}$) | 36,95 | 36,25 | 0,7 |
| | -7 ($10^{0.009}$) | neg | neg | | | -7 ($10^{0.1}$) | neg | neg | |
| | -8 ($10^{0.0009}$) | neg | neg | | | -8 ($10^{0.01}$) | neg | neg | |
| AHSV-7 | -2($10^{5.1}$) | 26,07 | 24,79 | 1,28 | AHSV-8 | -2($10^{4.6}$) | 23,9 | 23,75 | 0,15 |
| | -3 ($10^{4.1}$) | 28,16 | 28,09 | 0,07 | | -3 ($10^{3.6}$) | 27,38 | 27,02 | 0,36 |
| | -4 ($10^{3.1}$) | 31,72 | 30,79 | 0,93 | | -4 ($10^{2.6}$) | 32,18 | 31,92 | 0,26 |
| | -5 ($10^{2.1}$) | 36,48 | 35,09 | 1,39 | | -5 ($10^{1.6}$) | 34,01 | 35,27 | 1,26 |
| | -6 ($10^{1.1}$) | neg | 38,62 | | | -6 ($10^{0.6}$) | 38,32 | neg | |
| | -7 ($10^{0.1}$) | neg | neg | | | -7 ($10^{0.06}$) | neg | neg | |
| | -8 ($10^{0.01}$) | neg | neg | | | -8 ($10^{0.006}$) | neg | neg | |
| AHSV-9 | -2($10^{5.1}$) | 22,51 | 23,51 | 1 | | | | | |
| | -3 ($10^{4.1}$) | 26,35 | 25,67 | 0,68 | | | | | |
| | -4 ($10^{3.1}$) | 29,77 | 31,3 | 1,53 | | | | | |
| | -5 ($10^{2.1}$) | 32,07 | 34,89 | 2,82 | | | | | |
| | -6 ($10^{1.1}$) | 38,47 | 37,78 | 0,69 | | | | | |
| | -7 ($10^{0.1}$) | neg | neg | | | | | | |
| | -8($10^{0.01}$) | neg | neg | | | | | | |

**Cuantitative differences between replicates**

| | |
|---|---|
| **Mean Ct difference** | **0,90** |
| **SD** | **0,84** |

Rep., Technical replicate; Ct, cycle threshold; SD, standard deviation. Ct values obtained in each replicate and differences in absolute value between replicates are shown.
